# Highly accessible AU-rich regions in 3’ untranslated regions are hotspots for binding of proteins and miRNAs

**DOI:** 10.1101/042986

**Authors:** Mireya Plass, Simon H. Rasmussen, Anders Krogh

## Abstract

**Background:** MicroRNAs (miRNAs) are endogenous short non-coding RNAs involved in the regulation of gene expression at the post-transcriptional level typically by promoting destabilization or translational repression of target RNAs. Sometimes this regulation is absent or different, which likely is the result of interactions with other post-transcriptional factors, particularly RNA-binding proteins (RBPs). Despite the importance of the interactions between RBPs and miRNAs, little is known about how they affect post-transcriptional regulation in a global scale.

**Results:** In this study, we have analyzed CLIP datasets of 49 RBPs in HEK293 cells with the aim of understanding the interplay between RBPs and miRNAs in post-transcriptional regulation. Our results show that RBPs bind preferentially in conserved regulatory hotspots that frequently contain miRNA target sites. This organization facilitates the competition and cooperation among RBPs and the regulation of miRNA target site accessibility. In some cases RBP enrichment on target sites correlates with miRNA expression, suggesting coordination between the regulatory factors. However, in most cases, competition among factors is the most plausible interpretation of our data. Upon AGO2 knockdown, transcripts that contain such hotspots that overlap target sites of expressed miRNAs in 3’UTRs are significantly less up-regulated than transcripts without them, suggesting that RBP binding limits miRNA accessibility.

**Conclusions:** We show that RBP binding is concentrated in regulatory hotspots in 3’UTRs. The presence of these hotspots facilitates the interaction among post-transcriptional regulators, that interact or compete with each other under different conditions. These hotspots are enriched in genes with regulatory functions such as DNA binding and RNA binding. Taken together, our results suggest that hotspots are important regulatory regions that define an extra layer of auto-regulatory control of post-transcriptional regulation.

## Background

Post-transcriptional regulation controls gene expression at the RNA level and affects the properties and the amount of RNA from it is transcribed until it is translated into proteins. This regulation is performed mainly by RNA binding proteins (RBPs) and miRNAs, which primarily target the 3’ untranslated region (3’UTR) of transcripts. In animals, miRNAs usually function by promoting translational inhibition and decay of mRNAs[1]. In contrast, RBPs have a wider range of functions and are often involved in multiple post-transcriptional processes.

Although some miRNAs are predicted to target thousands of mRNAs[2], not all their predicted targets are down-regulated upon miRNA transfection[3], and many seem to be regulated only in certain cellular contexts or under stress conditions[4]. In some cases, miRNAs have even been found to promote translational activation[5] or increase mRNA levels. All these different complex functions suggest that miRNAs and RBPs take part in combinatorial regulation, where the *combination* of factors that bind to an RNA determines its fate.

In humans, more than 1000 RBPs[6, 7] and ~2500 miRNAs[8] are involved in this complex regulation. MiRNAs are known to act cooperatively to down-regulate mRNAs when bound close in space[3, 9, 10]. RBPs can compete for binding to AU-rich elements (AREs)[11] or cooperate in mRNA regulation[12, 13]. Moreover, they can compete and collaborate with miRNAs, or even perform opposite functions in different contexts[14]. For instance, AUF1 has been found to both compete with AGO2 for binding to the mRNA and cooperate with it[15]. Similarly, HuR has been found to compete with miRNA for binding[16, 17] but also to cooperate with miRNAs both to stabilize and degrade target mRNAs[18, 19].

Previous studies to understand the interactions between miRNAs and RBPs have been focusing either on single genes[20–24] or at the interactions between a single RBP and miRNAs using cross-linking immunoprecipitation coupled to high-throughput sequencing (CLIP-seq) data[13, 16, 17], and only recently it has started to be explored at a transcriptome-wide scale[25, 26]. In this work, we reanalyze a large collection of CLIP experiments from HEK293 cells to shed light on the complex interactions between RBPs and miRNAs. We demonstrate that post-transcriptional regulators bind preferentially in specific regions of 3’UTRs, which we call hotspots, where we find enrichment of both RBP and miRNA target sites. These hotspots, rather than being experimental artifacts as previously thought[26, 27], share characteristics of other regulatory elements: they are more accessible than other 3’UTR regions, more conserved, and enriched in AU-rich elements (AREs). Furthermore, in data from AGO2 knockdown (KD), we find that changes in the expression level of transcripts that contain free miRNA target sites are significantly different from those of transcripts that contain miRNA target sites within hotspots, which suggests that the RBPs binding in hotspots efficiently prevent RISC association and thus links hotspots directly to function. Interestingly, we observe that RBP hotspots are enriched in genes with roles in transcriptional and post-transcriptional regulation.

Taken together, these results suggest that post-transcriptional regulation is focused in hotspots within 3’UTRs that facilitate competition among regulators and modulate the functions of the regulatory network both at transcriptional and post-transcriptional level.

## Results

### RBP binding sites colocalize within 3’UTRs

To investigate the complex interactions between RBPs and miRNAs on a transcriptome-wide scale, we reanalyzed previously published CLIP data for 49 RBPs (Table S1) in HEK293 cells, which correspond to a total of 110 experiments. All the datasets were analyzed using the same pipeline in order to obtain a set of significant RBP binding sites that can be compared across experiments (see Methods). An analysis of the functions of these proteins using GO terms revealed that many are involved in similar processes, especially in post-transcriptional regulation of gene expression (Fig. 1a; Table S2). This is consistent with the observation that many of these RBPs show preferential binding on 3’UTRs, where most of these processes take place (Fig. 1b). On average, each mRNA expressed in HEK293 is bound by 14 RBPs in its 3’UTR, and 20% of them are bound by more than half of the RBPs analyzed. A detailed analysis of the distribution of RBP binding sites along 3’UTRs showed that most of them bind preferentially towards the 3’UTR edges, where there is also a higher density of miRNA target sites (Fig. 1c).

**Figure 1.**
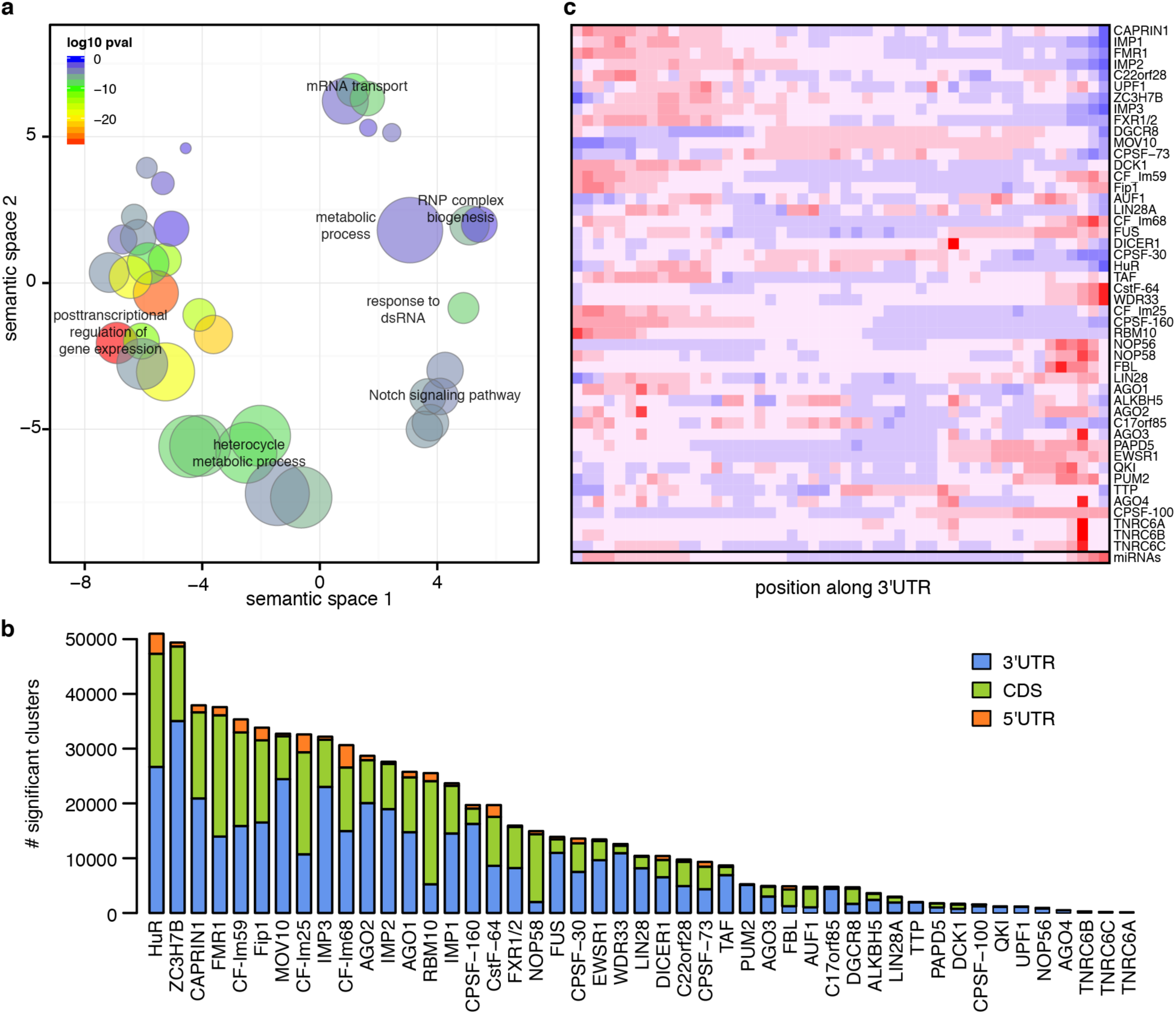
Summary of RBPs included in the analyses. **a)** Scatter plot summarizing the most significant GO-terms associated with the RBPs analyzed. Bubble sizes indicate the frequency of the GO-term in the GO database. The color of the bubbles represents how significant the term is in the set of RBPs analyzed. **b)** Distribution of significant clusters (FDR < 0.01) across gene regions. The height of the bars represents the amount of significant clusters in 3’UTRs (blue), coding region (green) and 5’UTR (orange). **c)** Heatmap showing the distribution of significant clusters across standardized 3’UTRs. The colors range from red (higher frequency) to blue (lower frequency).

After mapping the CLIP-seq data, clusters were identified and normalized by RNA-seq data to calculate an enrichment value of CLIP reads for each cluster. To understand if the observed positional bias indicates that the proteins bind in the same regions and therefore interact or compete with each other, we calculated spatial correlations between their cluster enrichments. For each pair of proteins, we calculated the Pearson correlation between CLIP enrichment values for each distance between ‐200 and 200 nt in each 3’UTR. The correlations calculated for these values were averaged over all 3’UTRs, giving us an average spatial correlation profile for the two RBPs. These correlations were then compared to those obtained from shuffling the clusters to calculate their z-scores at each position (Fig. 2a). In each row of Fig. 2a, the 401 nt windows around binding sites of an RBP are shown for each of the RBPs analyzed (columns). The higher the z-score, the more significant the correlation between two RBPs in a particular position is. In 746 out of 1084 pairwise combinations of RBPs (excluding pairing of an RBP with itself), we observed that the highest positional correlation z-score was in the +/-9 nt interval. This result reflects that for 69% of all RBP pairs, the most significant positional correlation was observed when the clusters of the two RBPs overlap in the same 3’UTR (Fig. 2a). If we consider only the RBP pairs that include AGO2, this percentage increases to 89% (Fig. 2a,b).

**Figure 2.**
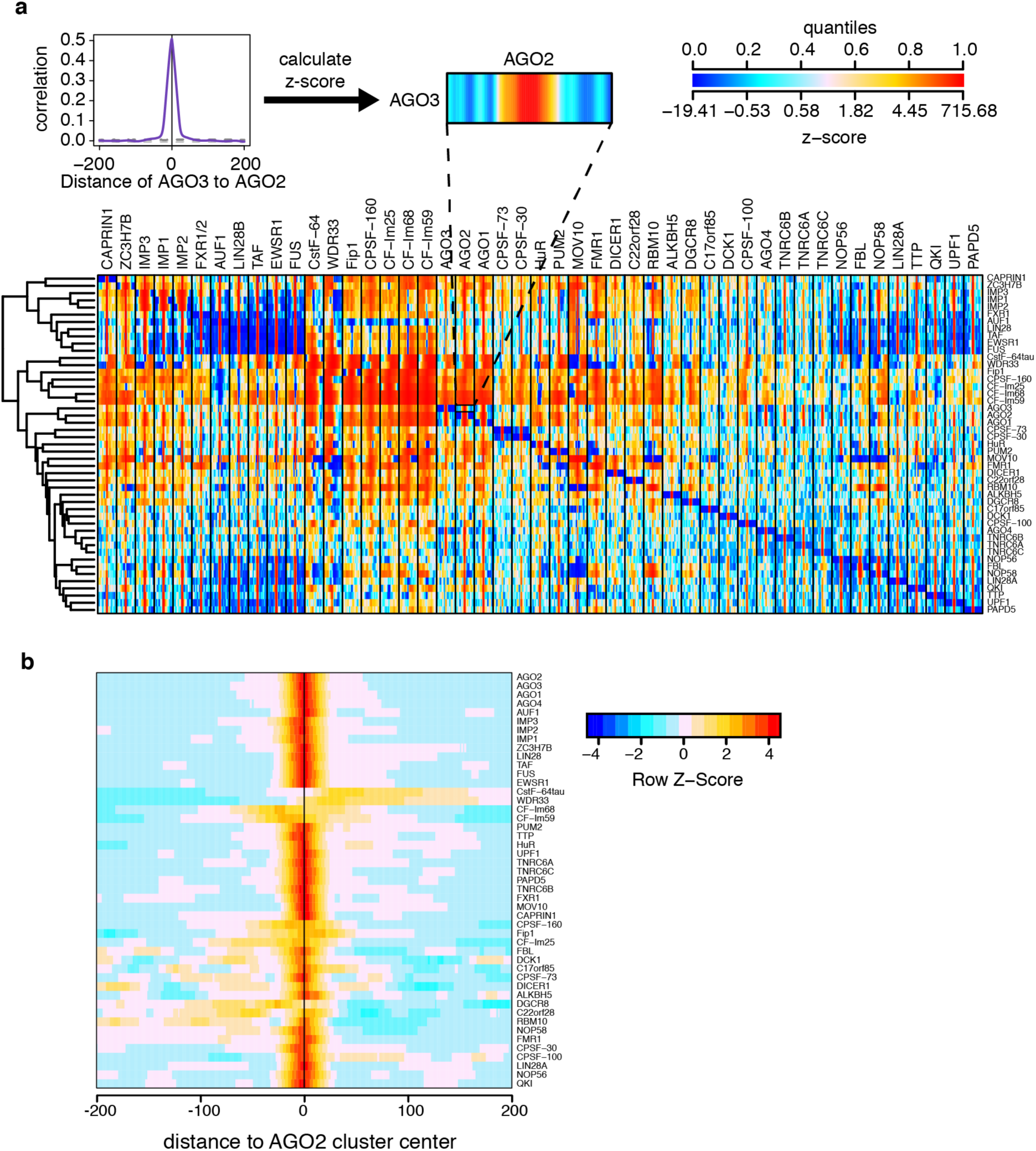
Positional correlation plots. **a)** Schematic representation of the z-score calculation for each RBP pair (top). Pearson correlation coefficients of cluster enrichments are calculated for each pair of proteins (purple line) in 401 nt windows and compared against the positional correlations obtained from shuffled clusters in the same region (grey lines show the top and bottom 5% random distributions). The random clusters are then used to calculate the z-score of the Pearson correlation coefficient at each position. The colors represent the z-scores of the correlations and range from blue (lower z-score) to red (higher z-score). All against all positional correlations calculated in this way are summarized in the heatmap (bottom). For each RBP in a column we show the strength of the positional binding each of the RBPs in the rows around its binding sites. **b)** Zoom on the positional correlations calculated as described before around AGO2 binding sites, row-normalized.

### RBPs preferentially bind on miRNA target sites

The previous analyses demonstrated that RBPs exhibit distinctive binding preferences around AGO2 binding sites. To further evaluate the correlations between RBPs and AGO2, we analyzed the enrichment distribution of their binding sites around predicted target sites of miRNAs expressed in HEK293 cells. As expected, the binding of AGO2 and the other AGO proteins peaked on top of predicted miRNA target sites, especially on target sites of miRNAs highly expressed in HEK293 cells (*hisites*) (Fig. 3a). Interestingly, a total of 26 out of 47 RBPs analyzed showed a significant enrichment around *hisites* (Fig. 3b and Fig. S1). In some cases, the enrichment profile peaked on miRNA target sites. In other cases, the enrichment increased across the miRNA target site, e.g. WDR33, or was less dependent on miRNA expression, such as in the case of HuR and EWSR1 (Fig. 3a and Fig. S1). Notably, the enrichment distribution around *hisites* in many cases resembles the positional correlation with AGO2 described before (Fig. 2b and Fig. 3b).

**Figure 3.**
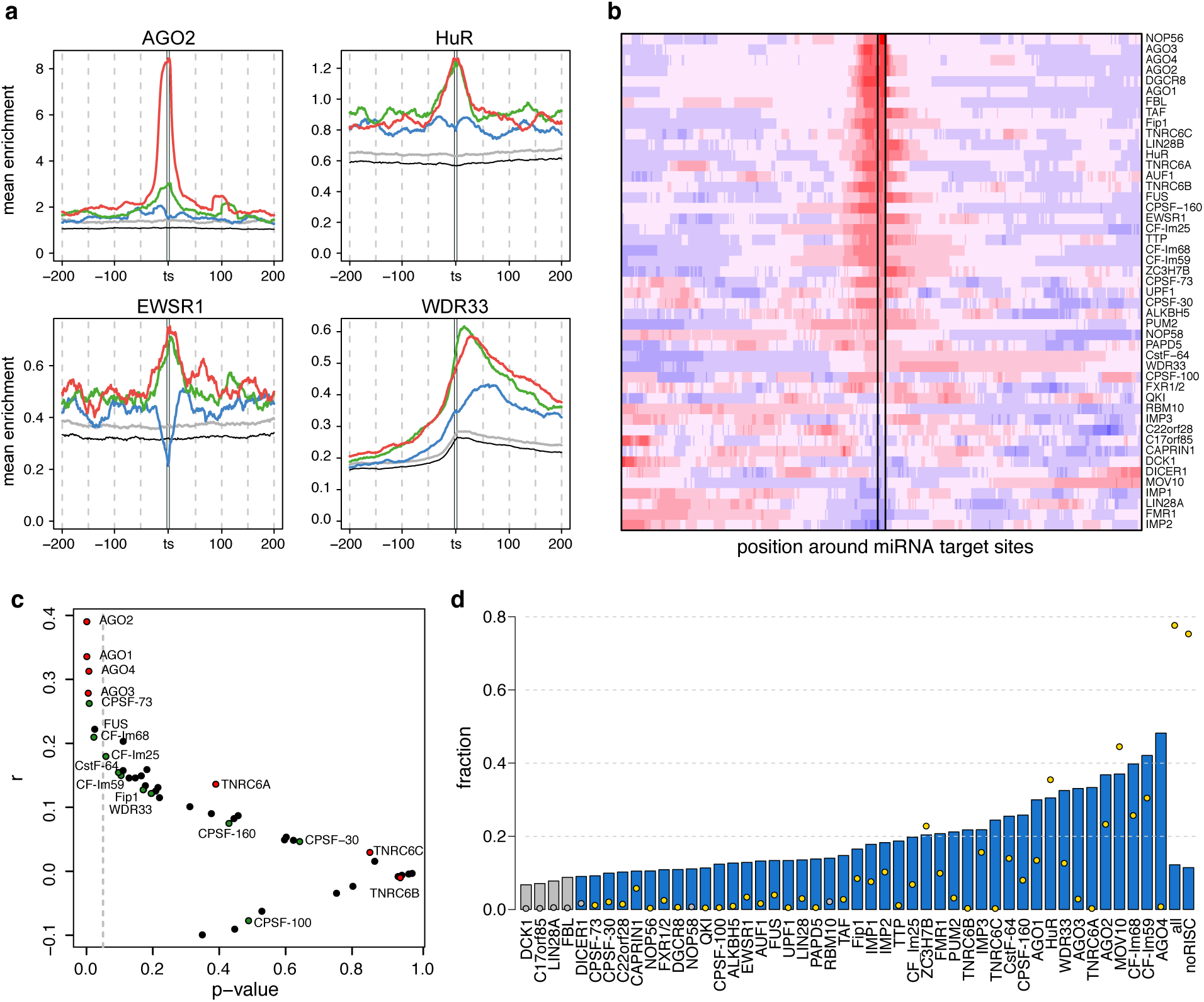
RBP enrichment around miRNA target sites. **a)** CLIP enrichment around miRNA target sites that are highly (red), moderately (green) and lowly (blue) expressed for AGO2, HuR, EWSR1 and WDR33 RBPs. The grey and the black lines show the maximum and the minimum enrichment values for the 90% confidence intervals around random miRNA target sites. **b)** Heatmap showing the distribution of CLIP enrichment values around *hisites*. The colors range from red (high enrichment) to blue (low enrichment). **c)** Scatter plot summarizing the correlation values between RBP enrichment on miRNA target sites and miRNA expression (y-axis) and their *p*-values (x-axis). **d)** Barplot summarizing the fraction of RBPs clusters on miRNA target sites (bar) and the fraction of *hisites* overlapped by RBPs (points). Colored bars (blue) and points (yellow) highlight the cases in which the fraction of RBPs or miRNA target sites is higher than expected by chance (empirical *p*-value < 0.01 using 100 random miRNA target site sets).

To gain insight into why RBPs have a strong positional bias around *hisites,* we calculated the correlation between RBP enrichment on target sites and miRNA expression level, i.e., the sum of expressions of all the miRNA targeting them. Our results show a significant correlation between RBP enrichment on target sites and miRNA expression not only for AGO proteins (AGO1-4), but also for the proteins from the polyadenylation complex CFIm68 and CPSF73 and FUS (Fig. 3c and Fig. S2).

### RBPs bind in regulatory hotspots containing miRNA target sites and AREs

For each RBP we also calculated the percentage of its clusters that overlapped *hisites*. This number ranges from less than 10% to more than 40% (Fig. 3d, bars), and usually is less than 5% of them (Fig. 3d, dots). Even though the overlap was small, in most cases the association between clusters and *hisites* was significantly higher than expected by chance (permutation test *p*-value < 0.01; colored bars and dots). Interestingly, the combined set of all RBPs overlapped more than 77% of *hisites* (75% excluding AGO and TNRC6 proteins), which suggests that the cumulative effects of all RBPs could have a crucial role in the modulation of miRNA function.

To evaluate the impact of all RBPs together, we looked at their binding in non-overlapping 50 nt windows across 3’UTRs. We observed that the windows with more than 3 RBPs binding were more frequent than expected if RBPs would bind independently (Fig. S3). We also observed that windows containing more RBPs displayed a stronger positional bias towards 3’UTR edges, similar to that of miRNA target sites (Fig. S4).

We performed several analyses on 3’UTR windows in order to identify the elements that are characteristic of these regulatory hotspots. The results from these analyses demonstrate that windows containing more RBPs are under stronger purifying selection. They are more conserved and have a lower sum of minor allele frequencies, which reflects both a lower frequency of SNPs and that the SNPs in the window are less frequent in the population (Fig. 4a).

**Figure 4.**
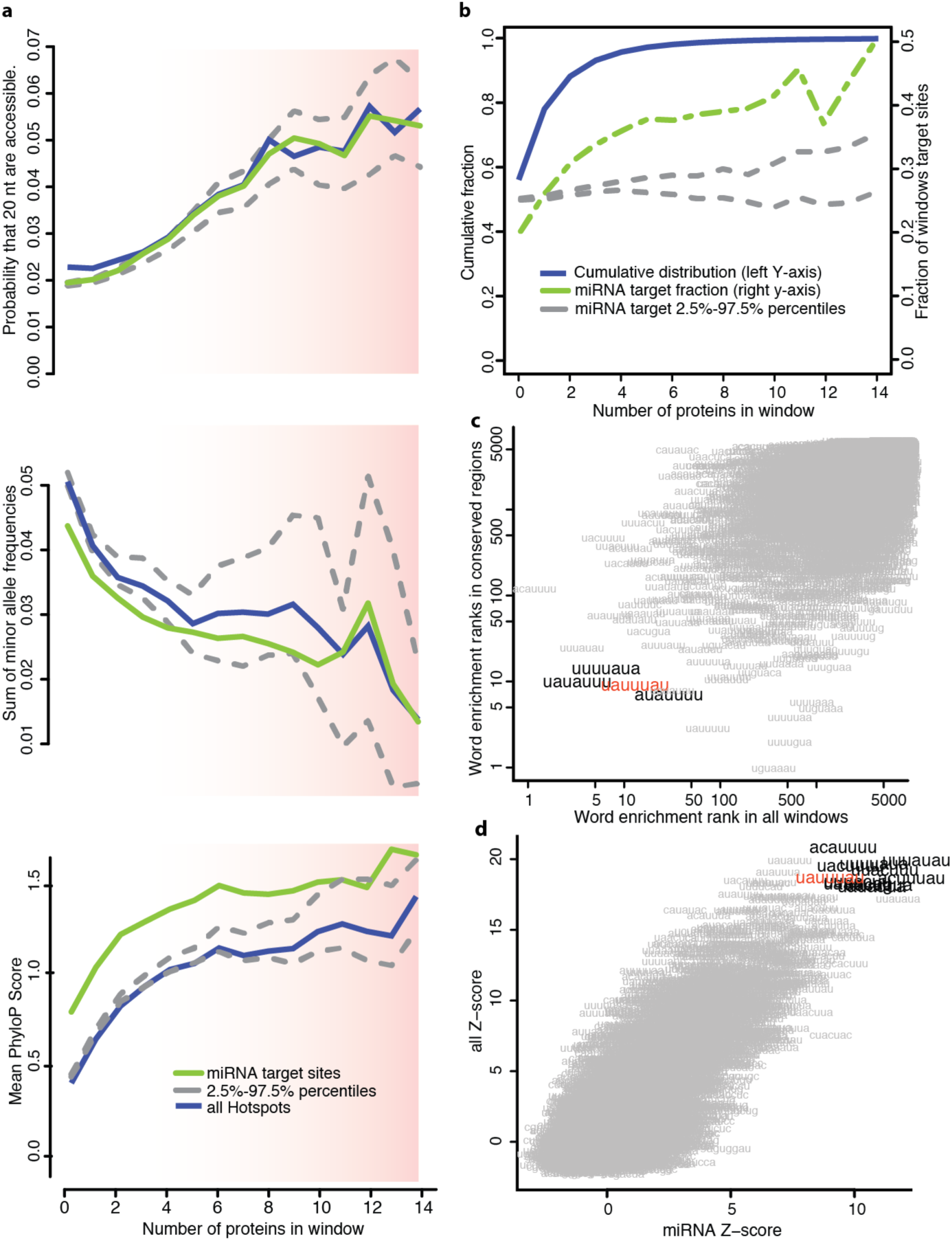
Characteristics of hotspots. **a)** Relation between accessibility (top), sum of minor allele frequencies (middle), and conservation (bottom) with the number of RBPs in a window for all windows (blue) and windows containing good predicted miRNA target sites (green). The grey dashed lines mark the 2.5% and 97.5% percentiles of the distribution obtained by generating 100 random miRNA datasets. **b)** Cumulative distribution function showing the fraction of windows (y-axis) bound by a number of RBPs (x-axis) for all windows (blue), windows containing miRNA target sites (dashed green line) and random miRNA target sites (grey lines). **c)** Scatter plot showing the correlation between the ranks of the 7-mers identified using cWords[28] ordering the windows according to mean PhyloP scores (y-axis) or the number of RBPs in a window (x-axis). Highlighted are the words that are in the top-20 in both analyses. The identified ARE UAUUUAU is highlighted in red. **d)** Scatter plot showing the correlation between the z-scores of the 7-mers identified ordering the windows according to the number of RBPs in all windows (y-axis) or only in windows containing miRNA target sites (x-axis). The words with the highest z-scores in both datasets (cut offs 17.5 and 8 for all and miRNA-containing windows respectively) are highlighted. In red is highlighted the ARE UAUUUAU.

Considering the dependencies among RBP binding sites, we decided to define hotspots in 3’UTRs as windows containing at least 5 RBPs. Using this definition, approximately 4% of all windows are classified as hotspots, whereas 56% of them are not bound by any RBPs (Fig. 4b). We noticed that the number of RBPs binding in a window is positively correlated with U-content (R= 0.21, *p*-value < 2.2e-16) and negatively correlated with G-content (R=-0.2; *p*-value < 2.2e-16) (Fig. S5). Additionally, windows targeted by several RBPs have much higher sequence accessibility, measured as the probability that at least 20 consecutive nt are unpaired (Fig. 4a).

We used cWords[28] to identify motifs enriched in hotspots. We identified several AREs, including UAUUUAU, among the top 20 ranked words enriched both in hotspots and in conserved regions (Fig. 4c). The core ARE element AUUUA was enriched in hotspots as well, although its frequency does not increase linearly with hotspot size (Fig. S5). We also noticed that the words enriched in hotspots overlapping miRNA target sites are very similar to those found in all hotspots (Fig. 4d, Table S3). Notably, we found an almost complete G-depletion in the top 100 words enriched in hotspots, which is consisted with the observation that hotspots have higher accessibility.

### RBPs compete at RBP hotspots

We have observed that hotspots are more conserved, more accessible, and enriched by miRNA target sites and AREs, including UAUUUAU, which has been associated with stronger miRNA effect and a mRNA stabilizing effect[3, 29]. Hence, we set out to assess the effect of RBP hotspots on *hisites* using previously published AGO2 KD microarray data[30]. For each transcript, we defined a new set of 50 nt windows centered on *hisites* and measured the effect of the presence of a hotspot (excluding AGO2 when defining the hotspots) overlapping 1, or 2 or more *hisites* in a transcript upon AGO2 KD. As a control, we used two sets of transcripts, one where all the *hisites* were in windows containing 2 or less RBPs and another one in which transcripts contained no *hisites* at all. By calculating the cumulative fractions of fold expression changes of transcripts upon AGO2 KD, we found that the presence of a hotspot overlapping *hisites* in a transcript prevents its upregulation upon AGO2 KD (two tailed KS test *p*-value = 0.0017 and 6.9e-08 for 1 and 2 or more target sites blocked compared to genes without hotspots on *hisites* respectively) (Fig. 5a). This result cannot be explained by differences in 3’UTR length or number of *hisites* in 3’UTRs (Fig. S6). Thus, it suggests that RBP hotspots can prevent the binding of RISC on miRNA target sites.

**Figure 5.**
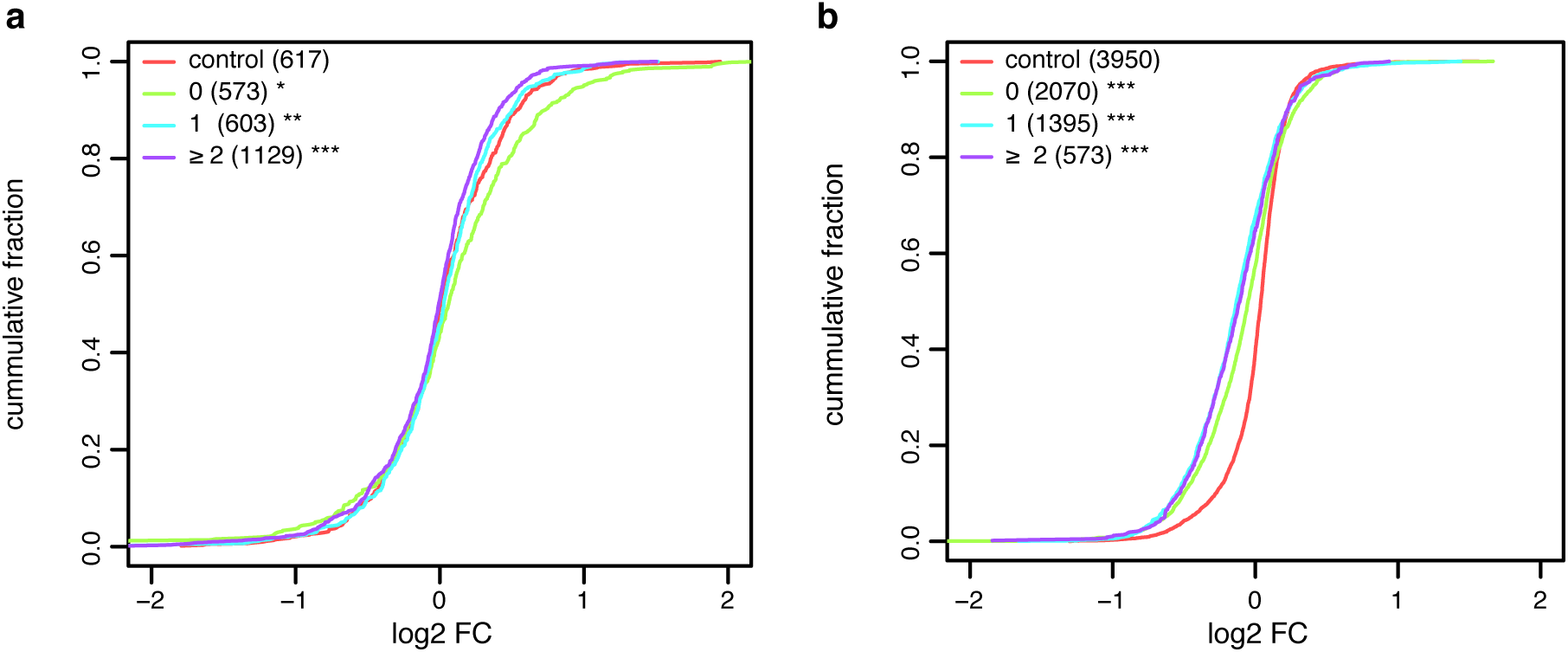
Competition within Hotspots. Cumulative fraction plot showing the effect of having 0, 1 or 2 or more **a)** *hisites* **b)** HuR binding sites overlapping RBP hotspots. As an additional control genes lacking **a)** *hisites* and **b)** HuR binding sites are shown. The x-axis shows the distribution of log2FC upon **a)** AGO2 or **b)** HuR KD. The y-axis shows the cumulative fraction of transcripts.

Additionally, we explored the function of hotspots overlapping the binding sites of other RBPs using published KD data. Similarly to the previous analysis, we defined hotspots centered on significant HuR clusters identified with CLIP data and measured the effect of the presence of a hotspot (without considering HuR) overlapping 1, or 2 or more HuR binding sites upon HuR KD[16]. As a control, we used transcripts where all the HuR binding sites were located in windows containing 2 or less RBPs (not counting HuR), or not bound by HuR. As expected, upon HuR KD, transcripts containing HuR binding sites were downregulated compared to transcripts not bound by HuR (Fig. 5b, two-tailed KS test *p*-value < 0.001 for transcripts with 1 and 2 or more HuR sites blocked compared to genes without hotspots on HuR sites). We also observed a small but significant difference between transcripts containing hotspots overlapping HuR binding sites and those that do not have them (KS test *p*-value < 0.001 in both cases). In this case, transcripts containing HuR sites overlapping hotspots are more downregulated upon HuR KD than transcripts containing HuR sites outside hotspots. This result suggests that upon HuR KD other RBPs with a negative effect on mRNA stability bind in those locations and thus promote mRNA downregulation. Accordingly, we found that 55% of the hotspots overlapping HuR sites contained AGO proteins, TNRC6 proteins, AUF1 or TTP, which are all known to be involved in promoting mRNA decay. Similar results were observed when analyzing the effect of hotspots overlapping AUF1 and TTP binding sites (Fig. S7). Taken together, these results show that RBP hotspots facilitate competition among RBPs and miRNAs, which affects post-transcriptional regulation.

### RBPs hotspots define a post-transcriptional regulatory network

In order to understand the biological function of RBP hotspots transcriptome-wide, we sought to characterize the transcripts containing hotspots. Transcripts with hotspots possess some features that suggest that they are under strong post-transcriptional regulation, as they have longer 3’UTRs (spearman correlation coefficient rho=0.17; **p*-value* = 1.2 e-60) while keeping approximately the same density of miRNA target sites (Fig. S8a,b). Furthermore, we also noticed that they are significantly higher expressed than transcripts without hotspots (rho=0.3; **p*-value* = 1e-191 Fig. S8c).

We used PANTHER[31] to characterize their functions using GO-terms. The most significant molecular function terms identified were polyA RNA binding, RNA binding and nucleic acid binding (*p-value* < 0.001; Table S4). Among the genes that contain these terms RNA binding proteins, splicing factors and transcription factors stand out (Fig. S9), which suggests that hotspots could be central in the regulation of both transcriptional and post-transcriptional processes.

## Discussion

In this work, we have analyzed a large collection of high-throughput experiments in order to better understand the complex interactions between RBPs and miRNAs in post-transcriptional regulation. Our results show that RBPs and miRNAs often bind in the same regions within 3’UTRs, which function as regulatory hotspots that facilitate competition between the regulators. These hotspots would therefore function in an analogous manner to promoter regions in accessible chromatin regions, and the RNA fate would depend on which of the regulators bind to the mRNA. In turn, this regulation would depend on external cues or post-translational modifications that modulate the competition between the regulatory factors. Interestingly, RBP hotspots are enriched in transcripts involved in transcriptional and post-transcriptional regulation, such as RNA binding proteins, splicing factors, transcription factors and translation factors (Fig. S9), thus suggesting that these regulatory hotspots play a role in an auto-regulatory network. This result is in agreement with recent findings that show that RNPs tend to regulate the mRNAs of other RNPs and themselves thus creating auto-regulatory networks in *Drosophila*[32].

We have used positional correlations to assess the interactions between RBPs assuming that RBPs that bind in the same location may interact. Surprisingly, we found that most RBPs (69% of all RBP pairs analyzed) tend to have overlapping binding sites (Fig. 2a). We confirmed some known positional correlations, such as those among polyA complex proteins[33], the IMP proteins, and the AGO proteins[34]. Moreover, we found correlations that were previously unknown. Some of these can be explained by the similarity in binding motifs, such as those among IMP proteins, HuR, TTP and AUF1[11, 13, 34]. However, the consistent correlation of all the RBPs analyzed with AGO2 had not been not previously described in literature. Additionally, the finding that many of these RBPs are also enriched on *hisites* further supported the positional correlations. It has to be noted that these miRNA target site predictions are independent of the CLIP data[35], which speaks against these overlaps being an artifact of the CLIP protocol.

One intriguing question is why the RBPs bind on miRNA target sites. If RISC directly interacts at miRNA target sites with a particular RBP, it would be expected that CLIP enrichment covaries with the expression of the miRNA that targets it. Yet, in most cases we did not find a clear correlation (Fig. S2) and thereby direct interaction is probably not the general mechanism to explain RBP enrichment at miRNA target sites. Nevertheless, we found a positive correlation for several proteins from the polyadenylation complex although it is only significant for CPSF73 and CFIm68 (Fig. 3c). This observation could indicate that miRNAs are involved in polyA-site selection. This hypothesis is in agreement with a recent publication that reported a significant overlap between AGO2 binding sites and m6A residues[36]. m6A residues have been related to the regulation of alternative polyadenylation, and thus, their overlap with AGO2 suggests a direct interaction between miRNAs and alternative polyadenylation.

We have also shown that RBP binding sites cluster in regulatory hotspots in 3’UTRs. These hotspots are more frequent than expected if RBPs would bind independently (Fig. S3) and are significantly enriched on predicted miRNA target sites (Fig. 4b). Furthermore, they are AU-rich (Fig. S5) and contain AREs, which are both conserved and overrepresented (Fig. 4c). AREs and AU-rich context of miRNA targets have previously been associated with effective miRNA target sites[3, 29, 37]. Therefore, our analyses suggest that RBP hotspots are functional regulatory elements in 3’UTRs.

Finally, we have shown that RBP hotspots regulate miRNA target site accessibility and favor the competition between miRNAs and RBPs in 3’UTRs. Upon AGO2 KD, transcripts containing miRNA target sites in hotspots are not significantly upregulated, which suggests that these target sites were protected by RBPs binding in the same hotspots (Fig. 5a). Interestingly, the opposite effect has been found upon KD of HuR, AUF1 and TTP. Upon KD of these RBPs, transcripts containing their binding sites in hotspots are more downregulated than those that have their binding sites isolated. A high fraction of those hotspots contain AGO2 or other down regulatory RBPs, which suggests that by removing the RBPs, other ones bind and affect mRNA stability (Fig. 5b and Fig. S7). These results are in agreement with recently reported findings that show that the presence of RBP binding sites of overlapping PUM1/2 or HuR binding sites reduce their impact on mRNA stability[26].

A previous study concluded that many regions found to be targeted by several different RBPs in CLIP-seq experiments are artifacts caused by biases in the experimental technique[27]. As a result, these regions have been excluded from previous works analyzing the combined effect of RBPs and miRNAs in post-transcriptional regulation[26]. However, for several reasons, we find that such background signal from CLIP cannot explain our conclusions. Firstly, it was described that background reads, i.e. the reads that appear in multiple datasets derived from a CLIP experiment of a control protein that does not bind RNA[27], are G-rich. In contrast, our RBP hotspots are characterized by a general G-depletion and are AU-rich (Fig. S5). Secondly, some of the analyzed RBPs do not show increased binding at the RBP hotspots (Fig. S10), indicating that the hotspots are not just a background phenomenon of the experimental method. Thirdly, the regulatory hotspots that we identify are experiencing increased selective pressure, as shown by the higher PhyloP scores and lower SNP frequencies (Fig. 4a), which support the conclusion that these regions are indeed functional regulatory elements and not background noise. Fourthly, we show that hotspots often coincide with predicted miRNA target sites, which are independent of CLIP, and it would be highly unlikely to happen if hotspots were artifacts. Besides, we see a clear functional effect of hotspots in the regulation of sequence accessibility both using KD data from AGO2 and other RBPs (Fig. 5 and Fig. S7). We believe that our stringent pipeline for the processing of the datasets, which includes duplicate removal, quality score aware mapping of reads, peak calling of clusters in transcripts, and normalization to RNA-seq, removes most of the reads that were shown to result in background when a less stringent data pipeline was used[27]. Accordingly, only 5% of our regulatory hotspots, i.e. windows containing 5 or more different RBPs, overlap background sites as previously defined[27]. Removal of these windows from our dataset did not alter the results reported in Fig. 4 (data not shown), which confirm our observation that RBP hotspots are not CLIP artifacts.

Many studies have previously investigated the interaction between miRNAs and RBPs using both experimental and computational methods[14, 25, 26, 38, 39]. Both competition and collaboration between miRNAs and RBPs have been described, but these interactions have been often portrayed as isolated events rather than a general mechanism in post-transcriptional regulation. In this work, we have shown that the overlap between miRNA target sites and RBPs is very extensive, with more than 75% of all *hisites* targeted by one or more of the RBPs analyzed (excluding AGO and TNRC6 proteins), thus suggesting that RBP hotspots play a major role in miRNA regulation and post-transcriptional regulation.

## Conclusions

Post-transcriptional regulation is one of the key processes involved in the regulation of mRNA levels and is in part controlled by the interaction between RBPs and miRNAs. With the development of high-throughput sequencing techniques, understanding the effects of these interactions and how they affect mRNA expression at a global scale has become possible. In this work, we have investigated the interactions in post-transcriptional regulation by integrating CLIP, RNA-seq and KD data in HEK293 cells. The results of our analyses show that RBPs and miRNAs target the same regions in 3’UTRs, which function as regulatory hotspots of post-transcriptional regulation. These hotspots are functional regions that coordinate post-transcriptional regulation. In them, we found not only evidence for competition among RISC and regulatory factors but also cooperation between polyadenylation complex proteins and RISC around miRNA target sites. Consequently, the outcome of the regulation is determined by the relative concentration of the effector factors and miRNAs in the cell, and thus more dependent on external cues that can modify the access of RBPs and miRNAs to the target mRNA.

Taken together, our analyses suggest that post-transcriptional regulation focuses in hotspots where trans-acting factors compete and cooperate. This organization would facilitate fast changes on mRNA expression induced as a response to environmental changes and facilitate cell adaptation to environment changes.

## Methods

### GO-term enrichment analysis

We obtained the significantly overrepresented biological process GO-terms associated with the RBPs included in the analysis using the gene enrichment analysis method performed by Panther[40]. The clustering and visualization of enriched GO-terms was done using REVIGO (http://revigo.irb.hr/)[41].

### Mapping and Processing of CLIP and RNA-seq datasets

115 CLIP (including CLIP-seq and PAR-CLIP) and 3 RNA-seq datasets were downloaded from GEO database[42]. The Sequence Read Archive (SRA) accession numbers of all the datasets analyzed can be found in Table S5.

Reads from all the experiments were preprocessed using custom python scripts. First, reads were trimmed to remove low quality scores and 3’ adapter sequences (only CLIP datasets). Next, we removed duplicates by collapsing all identical reads. After these steps, all reads longer than 19 nucleotides were further analyzed. Reads were mapped to the human genome (hg19) using bwa-pssm[43]. Then, all unmapped reads were then mapped to an exon-junction index containing all annotated unique exon-junctions from human Ensembl70 transcripts[44]. Only reads mapped at any of the steps with a posterior probability > 0.99 were considered for further analysis. For PAR-CLIP datasets, we used a custom matrix for scoring T to C mismatches assuming a 12.5% T to C conversion rate.

Datasets for the same proteins, or for different proteins with high correlation, were joined into a single dataset and analyzed together. Reads were clustered according to their genomic positions, requiring that at least 1nucleotide overlap. Significant clusters were calculated using Pyicos[45], using the exons from the longest protein coding transcript for calculating the randomizations. Only clusters with a false discovery rate (FDR) < 0.01 were considered for further analysis. The RNA-seq datasets were also joined and used together in further experiments. The statistics of the mapping and the datasets joined can be found in Table S5.

### Gene expression

For each 3’UTRs we calculated *m*_*k*_, the average number of RNA-seq base calls per nucleotide, normalized to *M*, the total amount of mapped RNA-seq reads in the experiment, as

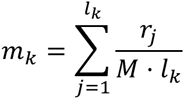

where *l*_*k*_ is the length of the 3’UTR for gene *k*, and *r*_*j*_ is the count of RNA-seq reads in position *j*.

### CLIP enrichment in 3’UTRs

For each transcript, we built a single-nucleotide resolution profile of the RBP binding sites, i.e. significant clusters with an FDR < 0.01 after peak calling, normalized to RNA-seq. The enrichment *e* of CLIP in a position *i* of a particular 3’UTR *k* is calculated as

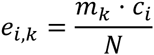

where *c*_*i*_ is the count of clip reads in position *i*, *N* is the total amount of uniquely mapped CLIP, and *m*_*k*_ is the average gene expression as defined above.

### Selection of miRNA target sites

Good miRNA target site predictions for conserved and non-conserved miRNAs were downloaded from miRNA.org (http://www.microRNA.org)[35]. Only target containing at least a 6-mer seed site were kept. Next, target sites were mapped to Ensembl70 transcripts[44]. Only target sites containing at least a 6-mer seed site included in the transcript were used.

We used small RNA-seq data (GSM1279922)[46] to estimate the expression levels of each miRNA. First, we selected from the dataset reads that were 15-27nt long, which corresponds to the length range of mature miRNAs. Next, we mapped the RNA-seq to a set of non-redundant human miRNA sequences downloaded from miRBase[8] using BWA-PSSM[43]. The expression of each miRNA was defined as the number of reads mapping to its mature miRNA sequence. We defined as expressed miRNAs only the top 20% of the mature miRNAs (155 miRNAs; minimum amount of mapped reads mapped 367).

We defined a set of non-overlapping target sites for the expressed miRNAs defined before. We overlapped the seed sites of their target sites and kept the one targeted by the most highly expressed miRNA. If several miRNAs shared the target site, we added their expression.

For some of the analyses we divided target sites according to their total expression, i.e. the sum of expressions of miRNAs targeting the same site, in three equally sized groups: highly expressed (*hisites*), moderately expressed and lowly expressed.

### Random miRNA target sites

To measure the significance of our results, we created 100 random sets of miRNA target sites containing as many target sites as the original set preserving their distribution along 3’UTRs. We divided the set of expressed genes with predicted miRNA target sites into 30 equal size groups with similar 3’UTR lengths. Then, for each target site in a particular 3’UTR, we assigned it to another of the 3’UTRs in the set. In case that the length of the new 3’UTR was different from that of the original 3’UTR, the relative coordinates of the target site were calculated so that it would have the same relative position within the 3’UTR in relation to its length. This procedure preserved the characteristic positional distribution of miRNA target sites along 3’UTRs.

### Mapping CLIP clusters on 3’UTR

The significant CLIP clusters for each of the RBPs were overlapped with the genes from Ensembl70[44] annotation using fjoin[47]. Only the longest protein-coding transcript for a gene was considered. If a cluster would overlap the CDS and a UTR region, the UTR annotation was assigned.

### 3’UTR positional data distribution

We analyzed the positional distribution of data across 3’UTRs of expressed genes (RNA-seq coverage >= 50%) and around *hisites*. Each 3’UTR was divided in 50 equally sized bins. For each bin, the mean value per nucleotide was calculated and then averaged across all expressed genes. In the case of CLIP data, the position of significant CLIP clusters (FDR < 0.01) was used to draw the profiles. In the case of hotspots, the position of the 50nt windows containing n (n=1,2…31) RBPs mapped on them was used. For miRNA target sites, the position of the target seeds in 3’UTRs was used.

### Positional correlation analysis

To find the positional correlation between the binding of two different proteins, we calculated the Pearson correlation between the enrichment values along a 3’UTR. If the value at position *i* is called for one RBP and *y*_*i*+*d*_ for the other RBP binding a distance *d* from the first, the Pearson correlation was calculated with fixed *d* over all positions *i*, in the interval from 1 to *I* – *d*, where *I* is the length of the 3’UTR (for negative *d*, the interval is from 1 – *d* to *I*). This was done for all values of *d* from ‒200 to 200. For each d, the correlation values were averaged over 3’UTRs. UTRs shorter than 400 were discarded.

The fluctuations of the correlation coefficients are heavily dependent on the number of CLIP sites. To estimate the background distribution, we shuffled the CLIP data in a way that preserved the clustering of tags. Clusters were defined as contiguous regions in which the enrichment value was above 10^-6^. The clusters identified in a sequence were moved to a random location in the sequence while ensuring at least one position in between clusters. After shuffling all sequences, positional correlations were calculated as above. This was repeated 100 times and for each *d*, the mean and standard deviation of the 100 values obtained in the shufflings were calculated. Using these estimates, the z-score was calculated for the unshuffled data. In Fig2a, the distribution of all z-scores calculated was considered and divided in 1000 quantiles. Each quantile was assigned a color from the scale, ranging from dark blue to red as shown. In Fig2b, z-scores were row-normalized and assigned a color using the same procedure as described above.

### Hotspot identification

To identify hotspots we divided the 3’UTRs of expressed genes (at least 50% RNA-seq coverage in the 3’UTR of the longest protein-coding transcript) in non-overlapping windows of 50 nt We overlapped the center of the RBP CLIP significant clusters with them and assigned each cluster to a single window. We also uniquely assigned each miRNA target site of expressed miRNAs in HEK293 cells to a window if the overlap between the seed site and the window was bigger than 5. Otherwise, the miRNA target sites were discarded.

### Simulation of RBP binding site distribution on hotspots

We simulated 10000 times the distribution of hotspot sizes by randomly sampling the binding location of the proteins assuming a uniform distribution of the RBPs in them. We considered the total amount of windows in which we observe significant clusters of each RBP and the total amount of windows in 3’UTRs (Table S6). The size distribution of hotspots from simulated and real data can be seen in Fig S3.

### Analysis of hotspot conservation

PhyloP scores[48] calculated from 100 vertebrate genome alignments (including hg19 human genome assembly) were downloaded from UCSC genome browser. For each of the 50nt nonoverlapping windows, we calculated the mean phyloP score across the window, discarding regions that were not present in any of the other species.

### Motif analysis

Word enrichment analyses were done using cWords[28]. The input data sets were made using the 3’UTR window data described above. For each window, we extracted its sequence and associated it to the number of RBPs binding in it. Using this method we defined two datasets: one containing all windows and another one containing only those overlapping target sites for expressed miRNAs.

In the first analysis, windows were ranked using the amount of RBPs binding in them. Thus, the resulting words were differentially enriched in windows according to the number of RBPs binding in them. In the second analysis, we ranked the windows using their mean phyloP score.

### RNA secondary structure accessibility of 3’UTR windows

We used RNAplFold[49] to calculated the sequence accessibility of the 3’UTRs. Specifically, we predicted the probability that 20 contiguous nucleotides in the sequence are unpaired using the parameters ‐u 20 ‐L 40 ‐W 120. The obtained accessibility values were then mapped to the 3’UTR windows and averaged across windows with the same number of RBPs binding and across windows with the same number of RBPs that overlap miRNA target sites.

### Minor allele frequency analysis

The complete data set of the 1000 genomes project containing all variants mapped to hg19 assembly[50] was downloaded. Of all the variants, we only used mutations regardless of their size and required them to be present in at least two individuals in a population of 5008. We calculated the mean of the sum of all minor alleles as 1 - major allele frequency regardless of which was the reference allele across windows as described above.

### Knockdown data analysis and processing

We downloaded the microarray data containing the expression values for AGO2 KD (GSM95818, GSM96819, GSM96816 and GSM96817)[30] and HuR KD (GSM738179, GSM738180, GSM738181, GSM738182, GSM738183)[16] from GEO database. We calculated differential expression upon AGO2 KD using the *limma* package[51] in R. We also downloaded processed data from KD experiments in AUF1[13] and TTP[52].

### Cumulative fraction plots

We defined 50 nt windows around *hisites* (35 nt upstream of the target site 3’ end, 14 nt downstream of the target site 3’end). If the windows extended beyond transcript boundaries, we shrank them so that they would fit inside the transcript. In each of these windows, we checked the presence or absence of each of the RBPs.

For each transcript we measured the amount of *hisites* that would be free, i.e. 2 or less RBPs (excluding AGO2) would bind in the window around the *hisite*, and the amount of hisites that would be blocked, i.e. 5 or more RBPs (excluding AGO2) would bind in the window around them. We used these measurements to divide the genes according to the amount of free or blocked *hisites* they contained in 3 groups: 0, where all target sites are free; 1, where only 1 target site was blocked; 2 or more, where 2 or more target sites were blocked. As an additional control, we added the rest of genes containing no *hisites*.

For cumulative fraction plots centered around RBP binding sites, we defined 50 nt windows centered around the binding sites of the RBP of interest. The groups of transcripts used to evaluate the rol of hotspots on RBP binding sites were built in an analogous manner to the one described above.

## Acknowledgements

We thank Anders Albrechtsen for help with the simulations in R. We also thank Jan Christiansen, Jeppe Vinther and Manuel Beltrán for helpful discussions and critical reading of the manuscript. This work was supported by the Carlsberg Foundation, the Novo Nordisk Foundation and the Danish Council for Strategic Research (Center for Computational and Applied Transcriptomics).

## Author Contributions

M.P., S.H.R. and A.K. conceived the project. M.P. and S.H.R designed and performed most of the analyses. A.K. performed positional correlation analyses. All authors contributed to writing the paper.

